# The Role of Frontal Eye Field in Saccadic Mixed-strategy Decision-making

**DOI:** 10.1101/2022.09.18.508403

**Authors:** Siwei Xie, Abdullahi Abunafeesa, Yong Gu, Mingpo Yang, Xiaochun Wang, Jiahao Tu, Dhushan Thevarajah, Michael Christopher Dorris

## Abstract

Game theory can predict the distribution of choices in aggregate during mixed-strategy games, yet the neural process mediating individual probabilistic choices remains poorly understood. Here, we examined the role of frontal eye field (FEF) in a decision-making task when macaques were trained to play a mixed-strategic game – Matching Pennies – against a computer opponent. Neuronal activities of FEF neurons predicted the animals’ upcoming saccadic choices and these activities became increasingly more selective as the choice deadline approached. Subthreshold electrical micro-stimulation applied in FEF also biased choices. Extended stimulation biased choices towards the preferred FEF vector whereas early termination of stimulation biased choices away from the preferred FEF vector. By contrast, micro-stimulation biased choices in the preferred direction during a non-strategic perceptual luminance discrimination task. We conclude that FEF is causally contributing to mixed-strategy decision-making process although the timing of FEF activation contributes to the decision process in a more non-linear manner during strategic compared to perceptual decision-making.

## Introduction

Mixed-strategy decision-making is deeply rooted in political, economic and interpersonal interactions where probabilistic actions are often required to avoid exploitation from opponents (a classic example being rock-paper-scissors). Game theory provides description on the proportional outcome where equilibria can be established among multiple rational agents in mixed-strategy games (i.e., 1/3 of each rock, paper and scissors). However, humans (and non-human primates) approach, but fall short of, the predicted equilibrium exhibiting a number of biases in their strategies (Worthy et al., 2012) (Lee et al., 2004). Therefore, to fully understand how choice strategies occur requires examination of the underlying neural mechanisms involved in monitoring choice outcomes and the selection of individual probabilistic choices.

Previous work has found neural correlates of strategic features during game play in multiple brain areas like prefrontal and parietal cortices (Coe et al., 2002; Parr et al., 2019; Seo et al., 2009). For example, latent values are encoded by prefrontal cortex (Barraclough et al., 2004) and lateral intraparietal cortex (Dorris and Glimcher, 2004) as well as abstract signals related to the historical sequence of game play, such as individual neurons representing the combination of choices and reward from previous trials (Seo et al., 2009; Seo and Lee, 2008, 2007). Whereas these cortical correlates are largely non-spatial/non-motor specific about “what could have been” (Lee and Dorris, 2014), the downstream midbrain area superior colliculus (SC) encodes the upcoming saccadic strategic choices directed to a specific saccadic vector. Neuronal activity becomes progressively more predictive for the upcoming saccade as the deadline to the action approaches. A disconnect remains in our understanding how these abstract historical and learning signals originating in the cortex are used to generate moment-to-moment actions in downstream structures. We hypothesize that the Frontal eye field (FEF) may play as critical bridge between these abstract cortical processes and more immediate pre-motor sub-cortical processes.

There are a number of reasons to suspect that FEF may be an important bridge between cortex and premotor SC in the selection of strategic saccades. First, FEF receives projections from multiple brain areas in prefrontal cortex and parietal cortex implicated in aspects of game play (Hutchison et al., 2012) including the dorsolateral prefrontal cortex (dlPFC), supplementary eye field (SEF), anterior cingulate cortex (ACC), lateral intraparietal area (LIP), (Barbas and Mesulam, 1981; Bates and Goldman-Rakic, 1993; Coe et al., 2002; Huerta et al., 1987). Plus, the topographic organizations of FEF (Bruce and Goldberg, 1985) provides a spatially organized platform where potential saccadic choices can compete during the selection process. Moreover, the frontal eye field sends controls signals to the SC through a dual pathway: an excitatory, direct projection to the intermediate layers of the SC (Wurtz et al., 2001), and an indirect pathway that disinhibits the otherwise inhibitory signals that the substantia nigra impart on the SC (Hikosaka et al., 2000). These connections can possibly provide guiding signals to the SC during strategic decision formation. Furthermore, neural correlates of evidence accumulation in perceptual decision-making process have been found in FEF (Ding and Gold, 2012), which could be tailored towards strategic decision-making.

To test the role of FEF in mixed-strategy decision-making, monkeys were trained to play the simple mixed-strategy game – Matching Pennies (Lee et al., 2004; vonNeumann and Morgenstern, 1944) – against a computer opponent. The activity of single neurons was recorded and multi-unit activity of neurons perturbed with electrical micro-stimulation to examine the correlational and causal role of FEF in selecting strategic saccades, respectively.

## Results

### Behavioral characterization

First, we examined how well monkeys competed against the computer opponent in the Mixed-Strategy game. We adapted the Mixed-Strategy game Matching Pennies (vonNeumann and Morgenstern, 1944) for an oculomotor setting (Fig. 1A). Monkeys were rewarded when they selected the same target chosen by a computer opponent. The computer’s algorithm (adapted from Lee et al. (2004)) attempted to minimize monkeys’ reward by exploiting statistical biases in monkeys’ patterns of choice or response to rewards (see Methods for details). Thus, although monkeys were free to choose, they would maximize averaged reward if they allocated their choices stochastically between the two targets from trial to trial. This stochastic strategy also corresponds with the Nash equilibrium for this game.

**Figure 1.**
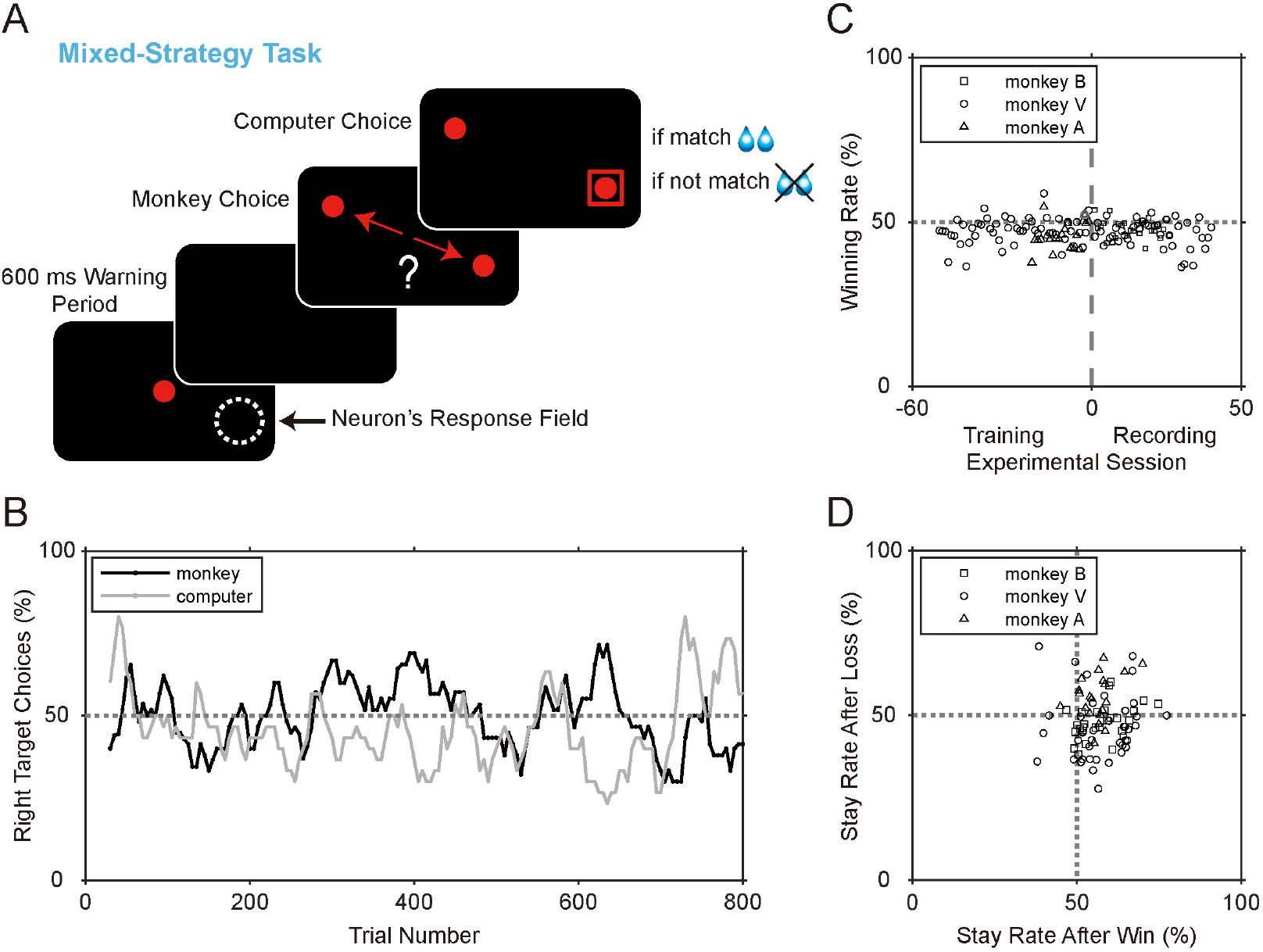
Behavioral summary. A) Schematic of Mixed-Strategy task. Each panel was presented to the monkeys chronologically. The white dashed circle represents the response field of a recorded neuron or end-point of the stimulation vector for a given FEF site. Red dots represent the visual targets presented on a black screen. The red square represents the choice of the computer opponent. Monkeys received liquid reward if their saccadic choice matched that of the computer opponent and no reward otherwise. B) The interplay between monkey’s (black) and computer’s (grey) rightward choices during an extended behavioral session. (Width of sliding window = 30 trials, window step = 5 trials). C) Average winning rate of 3 monkeys. Zero on the x-axis represents the beginning of neurophysiological recording sessions with training sessions beforehand. Data from Table 1. D) The likelihood that monkeys repeated choice direction (i.e., ‘stay’) after a rewarded choice (i.e., ‘win’) vs the likelihood of a stay after a losing choice during experimental blocks. Data from Table 1.

Broadly speaking, a dynamic competition between monkey and computer choices was observed that fluctuated around the mixed-strategy equilibrium of 50% within an individual experimental session (Fig. 1B). Overall, the probability of choosing the rightward target approximated 50% for all monkeys (Table 1). The overall reward rate provided a general measure of the predictability of the monkeys’ choices and their overall competitiveness in the mixed-strategy game. Their overall reward rate of 46.72% approached, but fell statistically short of, the 50% rate that would be obtained from a stochastic strategy (Table 1, Fig. 1C). Monkeys selected the same target as in the previous trial slightly more often (Table 1). Additionally, as has been reported previously (Lee et al., 2004), monkeys showed a Win-Stay Lose-Switch (WSLS) bias (Table 1; Fig. 1D)). Specifically, all three monkeys showed a WS bias while only one monkey showed a statistical LS bias (Table 1). Together, these results demonstrated that monkeys were competitive mixed-strategy game players, approaching the predicted Nash equilibrium. Like previous reports in humans (Worthy et al., 2012) and monkeys (Lee et al., 2004), they displayed subtle but statistically significant WSLS biases.

**Table 1.** Likelihood summary of monkeys’ choice behavior in Mixed-Strategy game

| Unit: % | P (right-ward) | P (win) | P (stay) | P (WSLS) | P (stay after win) | P (switch after loss) | P (stay after win) v.s.<br>P (stay after loss) | P (stay after win) v.s.<br>P (switch after loss) |
| --- | --- | --- | --- | --- | --- | --- | --- | --- |
| **Monkey B**<br>(N = 26 recording blocks) | 49.96 ±0.74<br>(p = 0.96) | 48.00 ±0.56<br>(p = 0.0013) | 52.48 ±1.02<br>(p = 0.022) | 54.38 ±0.61<br>(p = 2.1×10<sup>-7</sup>) | 57.28 ±1.40<br>(p = 2.3×10<sup>-5</sup>) | 51.81 ±1.05<br>(p = 0.096) | (p = 3.3×10<sup>-7</sup>) | (p = 0.0015) |
| **Monkey V**<br>(N = 40 recording blocks) | 47.42 ±1.48<br>(p = 0.086) | 46.46 ±0.65<br>(p = 3.1×10<sup>-6</sup>) | 51.60 ±1.09<br>(p = 0.15) | 55.42 ±1.08<br>(p = 1.2×10<sup>-5</sup>) | 57.67 ±1.42<br>(p = 3.7×10<sup>-6</sup>) | 53.86 ±1.56<br>(p = 0.018) | (p = 1.9×10<sup>-6</sup>) | (p = 0.085) |
| **Monkey A**<br>(N = 20 training blocks) | 49.37 ±0.94<br>(p = 0.51) | 45.58 ±0.89<br>(p = 8.5×10<sup>-5</sup>) | 55.90 ±1.14<br>(p = 5.3×10<sup>-5</sup>) | 49.61 ±0.89<br>(p = 0.66) | 56.16 ±1.18<br>(p = 4.8×10<sup>-5</sup>) | 44.43 ±1.54<br>(p = 0.0019) | (p = 0.71) | (p = 4.4×10<sup>-5</sup>) |
| **In total**<br>(N = 86) | 48.64 ±0.75<br>(p = 0.074) | 46.72 ±0.41<br>(p = 6.0×10<sup>-12</sup>) | 52.87 ±0.67<br>(p = 4.8×10<sup>-5</sup>) | 53.76 ±0.62<br>(p = 4.1×10<sup>-8</sup>) | 57.20 ±0.83<br>(p = 2.1×10<sup>-13</sup>) |  | (p = 7.6×10<sup>-10</sup>) | (p = 1.2×10<sup>-5</sup>) |

All cells refer to Mean ±SEM. Two-tailed one-sample t-test against 50% for columns showing simple likelihoods, but two-tailed paired t-tests for last two columns comparing likelihoods

For the analyses that follows, it is not critical that monkeys reach the Nash equilibrium precisely, nor be aware that they are competing against another agent. Rather, for our purposes, the matching pennies game elicited voluntary and largely unpredictable choices from trial-to-trial that allowed us to examine underlying generative stochastic process of strategic decision at the neural level.

### Neuronal activity was selective for upcoming choice

A total of 77 FEF neurons were recorded (monkey B: N = 35; monkey V: N = 42), 48 of which satisfied our neuronal criteria while the monkey’s behavior was also sufficiently reliable for inclusion in our data set (monkey B: N = 26; monkey V: N = 22; see Methods for details). We recorded neuronal responses during the Mixed-Strategy task (blue; monkey B: 26/26; monkey V: 20/22), Unpredictable task (green; monkey B: 18/26; monkey V: 18/22), and Predictable task (red; monkey B: 10/26; monkey V: 6/22), respectively. The number of neurons diminished in successive blocks either due to degraded isolation or diminished behavioral performance (see Methods for details). All the targeted neurophysiological sties were at the anterior bank of the left arcuate sulcus (Fig. 2A).

**Figure 2.**
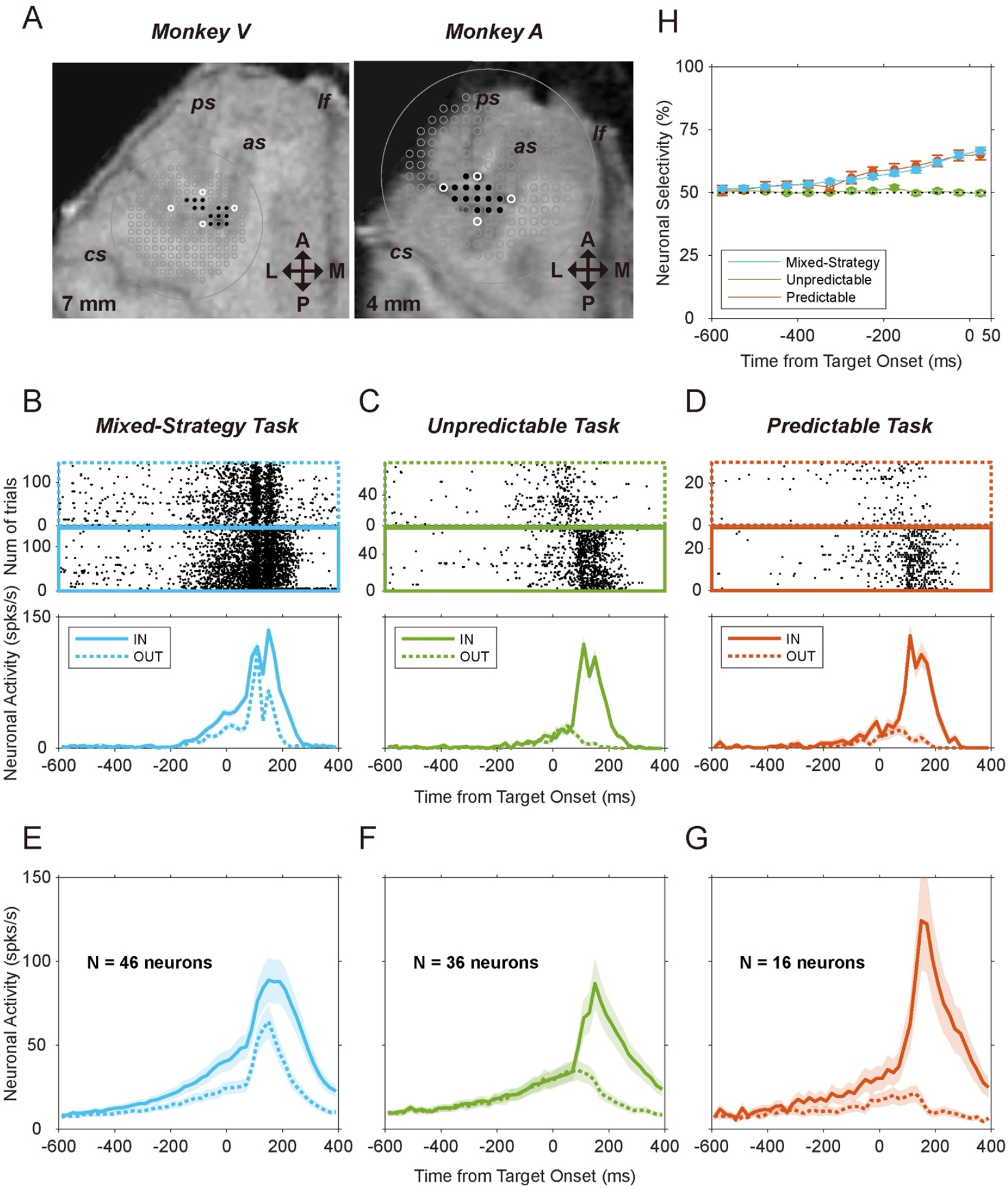
Summary of neuronal recording experiments. A) Electrode positioning grids are overlaid on brain MRI scans from 2 monkeys. Although difficult to visualize here, grids and associated electrode penetrations are tilted 25° to the left of the midline in the coronal plane. Arrows labeled A and P indicate the directions of true anterior and posterior axis. Arrows labeled M and L reflect the trend of medial and lateral axis, with 25° angle to the horizontal plane. The white circles represent the location of reference electrodes placed in the grid during MRI scans. Black dots indicate grid locations from which neurophysiological data was collected. The abbreviations of ps, as, lf, and cs indicate principal sulcus, arcuate sulcus, longitudinal fissure, and central sulcus, respectively. B-D) Activities of a representative FEF neuron during B) Mixed-Strategy (blue), C) Unpredictable (green), and D) Predictable (red) tasks. Raster plots (top panels) and post-stimulus-time-histograms (PSTH – bottom panels) are sorted based on monkeys’ choices towards (IN – solid lines) or away from (OUT – dashed lines) the response field target. E-G) Mean population activities during E) Mixed-Strategy, F) Unpredictable, and G) Predictable tasks. Shaded regions represent the ±SEM. H) Evolution of neuronal selectivity over time. Receiver operating characteristic (ROC) analysis for IN vs OUT neuronal activity during Mixed-Strategy, Unpredictable, and Predictable tasks. Same data sets as panels E-G, respectively. Error bars represent ±SEM. The filled points represent selectivity that was significantly greater than chance level (dashed line) (permutation test, 1000 permutations, p< 0.05).

We searched for neuronal correlates of strategic decision formation during the 600 ms warning period because monkeys initiated their choice shortly after the two targets were presented. Offline, we could then dissociate activities associated with neuron’s preferred choices (IN – solid lines) and neuron’s non-preferred choices (OUT – dashed lines) respectively (Fig. 2B-G).

As shown from a representative neuron (Fig. 2B), neuronal activity gradually increased before both preferred and non-preferred choices (see Methods) but rose to a higher level preceding the preferred choices. Consistent with the example neuron, average population activity also rose for both strategies with increased selectivity throughout the warning period (Fig. 2E, Table 2).

**Table 2.**
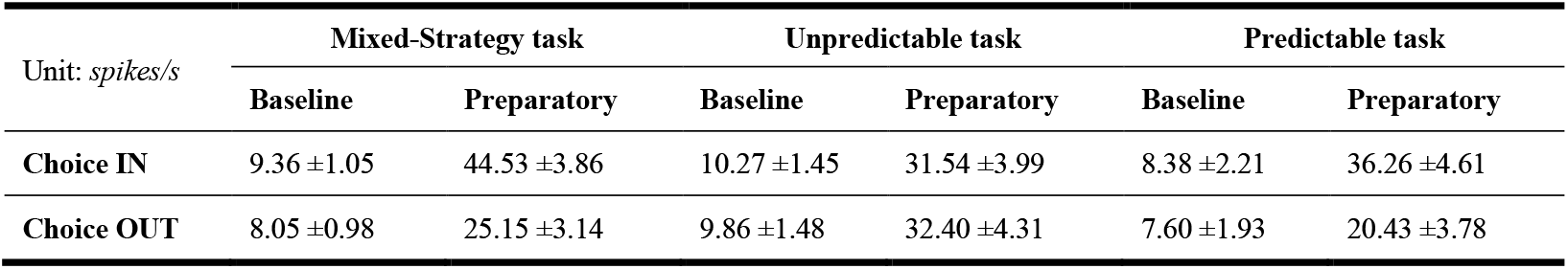
Averaged epoch activities during three tasks

| Unit: *spikes/s* | Mixed-Strategy task |  | Unpredictable task |  | Predictable task |  |
| --- | --- | --- | --- | --- | --- | --- |
|  | Baseline | Preparatory | Baseline | Preparatory | Baseline | Preparatory |
| **Choice IN** | 9.36 ±1.05 | 44.53 ±3.86 | 10.27 ±1.45 | 31.54 ±3.99 | 8.38 ±2.21 | 36.26 ±4.61 |
| **Choice OUT** | 8.05 ±0.98 | 25.15 ±3.14 | 9.86 ±1.48 | 32.40 ±4.31 | 7.60 ±1.93 | 20.43 ±3.78 |

The final preparatory activity (0-50 ms after the visual target point appeared) was significantly higher when the neuron’s preferred saccadic choices were made compared to its non-preferred choices (Fig. 2E, Table 2, paired t-test, p = 2.3×10^−11^). An ideal observer who used only the distributions of neuronal activities could predict saccadic choices with increasing reliability (Neuronal selectivity, Fig. 2H, blue) suggesting a gradual selection process.

### Control Experiments for Neuronal Recordings

The Predictable task and Unpredictable tasks allowed us to examine how FEF neurons responded under comparable non-strategic conditions that did or did not allow advanced saccade planning, respectively (Fig. S1, see Methods). Warning period activity in Unpredictable task, when upcoming saccades timing was certain but direction was uncertain, increased for both preferred and non-preferred choices (Table 2, paired t-test, non-preferred: p = 6.1×10^−6^; preferred: 3.9×10^−6^). However, this preparatory activity did not select between two targets (Table 2, paired t-test, p = 0.38) remaining near chance level (50%) throughout (Fig. 2H, green).

In contrast, preparatory activity in the Predictable task, when upcoming choice could be planned with both timing and location certainty, was highly selective for preferred vs non-preferred saccades (Fig. 2H, red). It is worth noting that even for non-preferred saccades, FEF activity increased throughout the warning period (Table 2) which may reflect responses to saccadic timing (Reddi and Carpenter, 2000).

Together, these results demonstrated that FEF activity was predictive of monkeys’ saccadic choices that could be planned voluntarily or in advance. To determine if FEF plays more than a correlative role in choosing strategic saccades we next attempted to artificially bias choices with electrical micro-stimulation.

### Artificially manipulating strategic choices

Up to this point, our neurophysiological results are purely correlational. These results cannot distinguish whether FEF is actively involved in the strategic decision-making process or whether it passively reflects ongoing decision processes being conducted elsewhere. To test whether FEF plays a causal role in strategic decision-making, we applied subthreshold micro-stimulation to the FEF. If this changed the likelihood of choices towards the two targets this would indicate that FEF is actively involved in selecting mixed-strategy saccades.

Importantly, our goal was to use low level electrical micro-stimulation to bias the strategic decision process (see Methods for details) rather than directly (and trivially) to elicit saccades with supra-threshold FEF stimulation (Bruce et al., 1985). Only experiments in which no saccades were triggered by ongoing micro-stimulation were included in the results that follow (Fig. S3).

As closely as possible, we replicated the protocols of Thevarajah et al. (2009) which examined the role of the SC in strategic decision-making so we could directly compare SC functioning to that of the FEF (see Discussion). Micro-stimulation was applied only during the first 500 ms of the warning period (MSstim500). Hence, micro-stimulation was terminated 100 ms before target presentation and more than 250 ms (Monkey B: 270 ms ±4 ms; Monkey V: 291 ms ±1 ms; Monkey A: 300 ms ±2 ms; Median ±SEM) before the onset of the saccadic choices. Micro-stimulation of FEF reliably biased choices (Fig. 3B, paired t-test, N = 103, p = 3.2×10^−13^), suggesting that FEF indeed was actively involved in the saccadic strategic decision-making process. But to our surprise, subthreshold micro-stimulation during first 500 ms of warning period biased strategic choices *away* from the supra-threshold stimulation vector.

**Figure 3.**
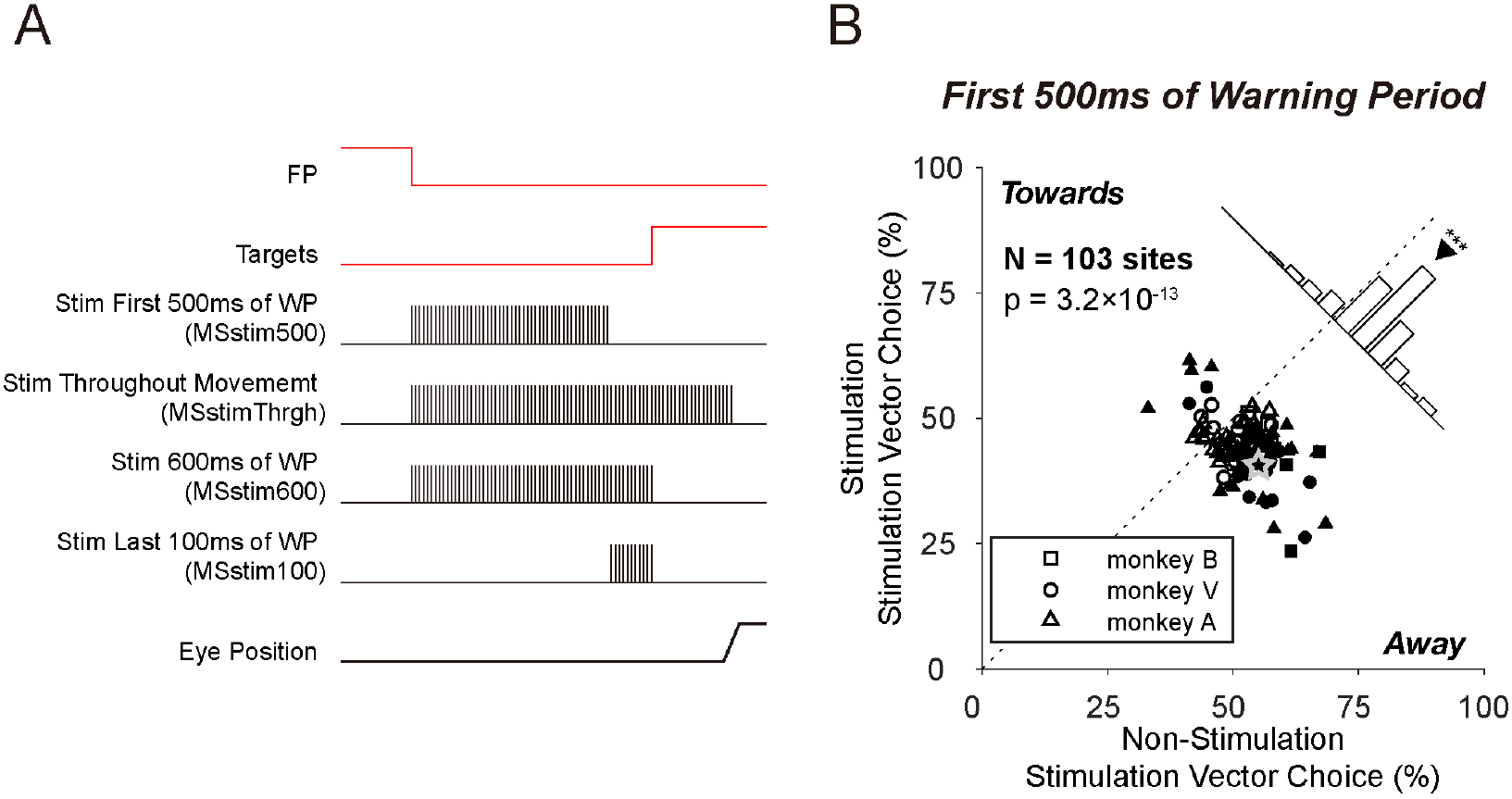
Perturbing strategic choice formation with FEF subthreshold electrical micro-stimulation. A) Schematic illustrating the various timings during which micro-stimulation was applied (vertical rasters). In each block, subthreshold micro-stimulation (0.25 ms biphasic pulses, 100 Hz, site specific intensity (see Methods)) was applied randomly on half of the trials. B) Results from the stimulation of the first 500 ms of the warning period (Stim First 500ms of WP). Percentage of choices to the stimulation vector target during stimulation versus the non-stimulation trials. The diagonal line represents unity. Filled symbols indicate statistical significance within an experimental session (Chi-square test, p< 0.05). The histogram displays the distribution of the difference between the x and y axes across the all experiments, and the downward-facing triangle denotes the mean (bin width = 6.6%). (*** denotes paired t-test, p< 0.001, and none for not significant). The grey star denotes the example site shown in Fig. S3.

It is unclear why applying micro-stimulation to the FEF during the warning period biased choices away from the stimulation site vector. One reason could due to withdrawing the stimulation 100 ms before target presentation and over 250 ms before the saccade itself. In fact, a more standard protocol applying stimulation on FEF during other cognitive tasks was durative until saccade execution (Burman and Bruce, 1997; Ditterich et al., 2003). Therefore, we tested the effect of micro-stimulation throughout the warning period and until the end of the eye movement (MSstimThrgh). Indeed, using these more traditional parameters, subthreshold stimulation throughout movement reliably biased strategic choices towards the stimulation vector (Fig. 4A, paired t-test, N = 94, p = 9.6×10^−15^).

**Figure 4.**
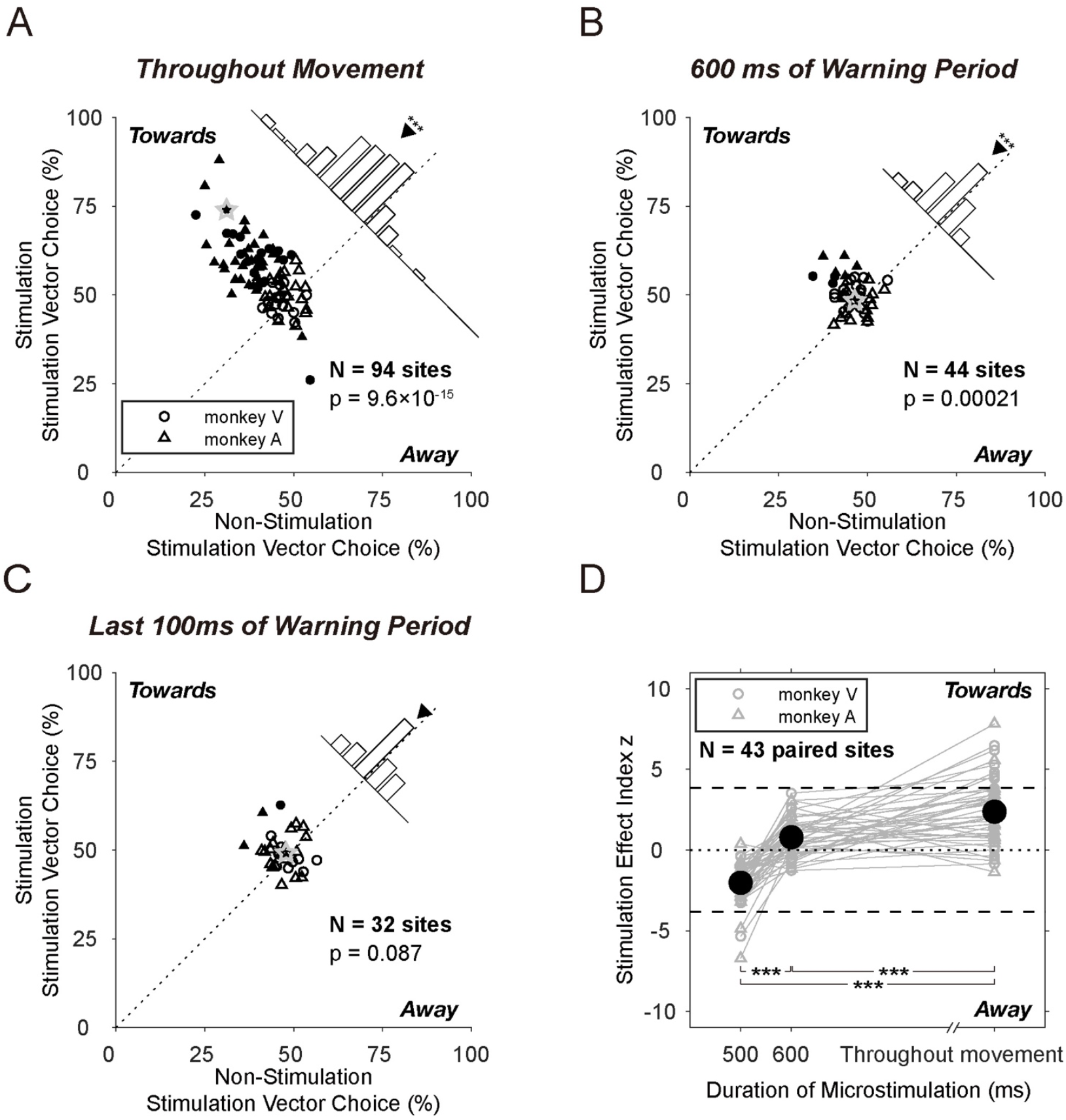
Effect of stimulation timing on strategic decision formation. A-C) Percentage of choices to the stimulation site target during stimulation versus the non-stimulation trials. Stimulation was applied for A) throughout the warning period until the saccade was detected online (N = 94 sites), B) throughout the 600 ms warning period (N = 44 sites), and C) during the last 100 ms of the warning period (N = 32 sites). Format the same as Fig. 3B. D) A comparison of stimulation effects across all 3 timing blocks (N = 43 sites). Data from shared sites are connected with grey lines. The horizontal dashed lines represent the level of statistical significance for individual experiments (Chi-square test, p = 0.05). Brackets with asterisks show the significance of population difference (paired t-test, *** for p< 0.001).

One reason stimulation throughout movement could bias choices towards the stimulation vector was that stimulation affected the visual or motor processing after target presentation rather than earlier decision processes during the warning period. To test this possibility, we applied micro-stimulation throughout the warning period and terminated it simultaneous with target presentation (Fig. 3A, MSstim600). Stimulation throughout the 600 ms warning period biased monkeys’ strategic choices towards stimulation vectors (Fig. 4B, paired t-test, N = 44, p = 0.00021) suggesting that this biasing effect is not reliant on perturbation of sensory or motor processes.

The effect of stimulation timing on bias direction was a consistent phenomenon across stimulation sites. Fig. 4D shows the progression of saccade direction from away to towards the stimulation site as stimulation was extended for a subset of 43 sites in which multiple duration stimulation experiments were conducted.

Therefore, it appears that stimulation at the very end of the warning period seemed to be of particular importance for biasing the strategic decision process. Indeed, subthreshold micro-stimulation applied only during the last 100 ms of the warning period had a tendency – but not did not reach statistical significance – to bias strategic choices towards the stimulation vectors (paired t-test, N = 32, p = 0.087; Fig. 4C, MSstim100).

Perturbing only the last 100 ms at the end of the warning period was not as effective as extending the stimulation from 500 ms to 600 ms (paired t-test for N = 32 sites on which all 3 stimulation conditions were applied, p = 9.9×10^−8^; bootstrap for all sites across conditions, iteration = 100, paired t-test for generated distributions, p = 7.0×10^−12^). In other words, merely stimulating at the late decision epoch was not equivalent to extended stimulation leading up to this period, meaning that the perturbation effect was non-additive, or more specifically, accumulative along the dimension of time, which suggests a non-linear interplay of timing and the strategic decision process.

The effect of micro-stimulation on saccade reaction time (SRT) for preferred and non-preferred saccades was quantified respectively (Fig. S5) using multiple regression analyses (see Methods). Overall, stimulation had a mild effect on SRTs in absolute terms (i.e., < 10 ms change in most cases) although statistically significant in some cases (Fig. S5 asterisks). Micro-stimulation induced SRT differences were consistent with the biasing effect on choices, i.e., lengthening SRT for saccades whose proportions were reduced by stimulation, and shortening SRTs for saccades whose proportions were increased by stimulation.

Altogether, our experimental results show that FEF is causally involved in the saccadic strategic decision-making process. However, early termination of activation biased strategic choices away from the preferred FEF vector whereas extended and later activation biased choices towards the preferred FEF vector. Hence, timing of FEF activation was an important factor as to perturbation effect, which appeared to be non-additive through the late epoch of saccadic strategic decision.

### Effects of subthreshold micro-stimulation in a perceptual decision task

Previous studies involving FEF micro-stimulation improved cognitive functions such as spatial attention and perceptual decision-making (Burman and Bruce, 1997; Moore and Armstrong, 2003; Moore and Fallah, 2001; Schafer and Moore, 2007) at the preferred location. These studies differed from ours in two ways. First, our initial perturbation schedule ended before presentation of sensory targets and the motor act. Indeed, above we demonstrated that extending the perturbation promoted choices towards the preferred vector. Second, these studies involved the participation of FEF in perceptual rather than strategic decision framework. To test whether there was a fundamental difference between how FEF contributes to perceptual and strategic decision-making, micro-stimulation was applied to FEF using the same parameters but applied to a more standard perceptual task.

We designed a perceptual decision-making task (Luminance Discrimination task, Fig. 5A) that closely mimicked the parameters of the Mixed-Strategy task. Subthreshold micro-stimulation was applied to the FEF for the first 500 ms of the displaying period. Rather than a strategic choice, monkeys were tasked with making a perceptual judgement concerning which of two visual stimuli were brighter.

**Figure 5.**
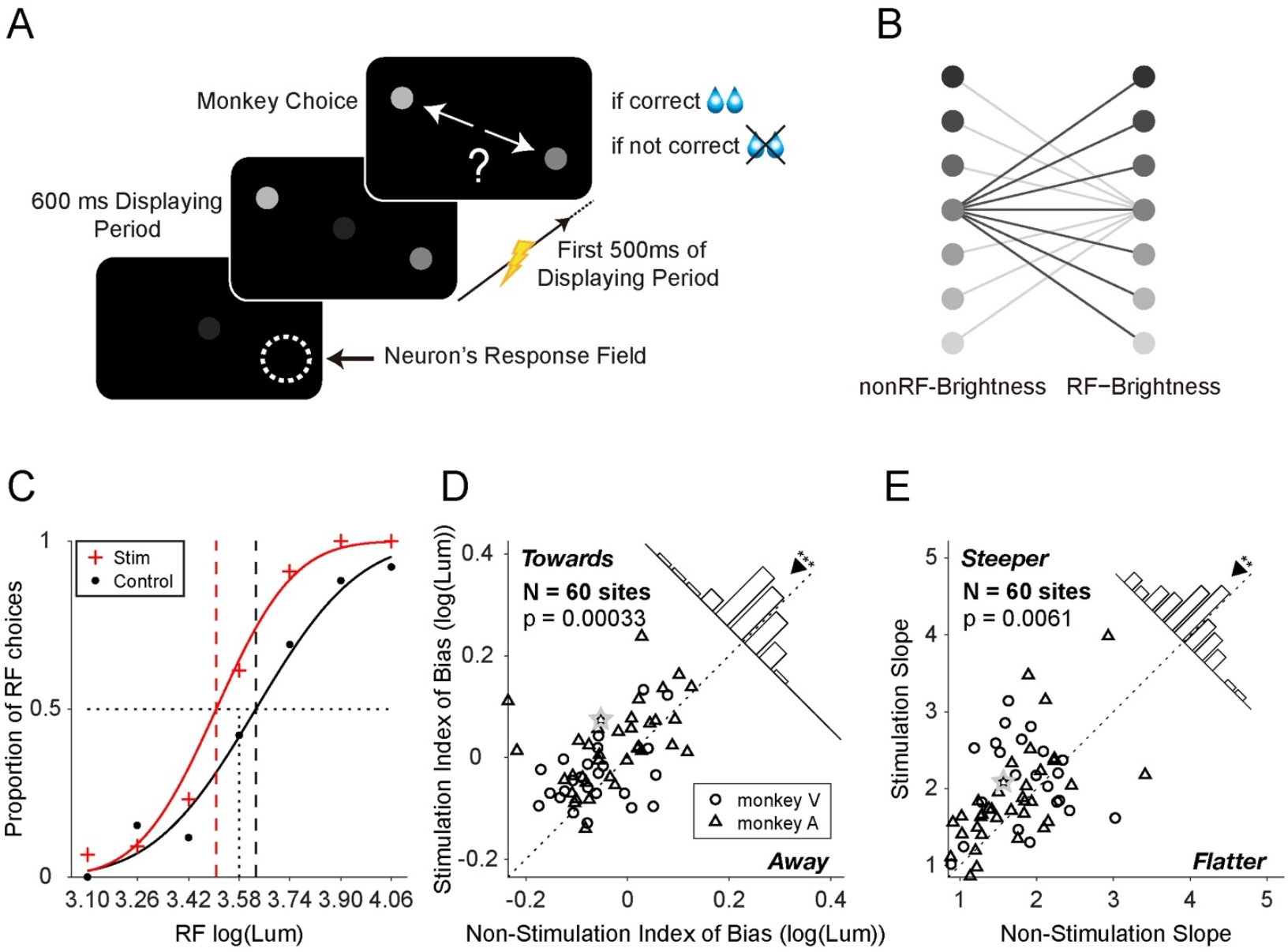
Subthreshold micro-stimulation biases choices during Luminance Discrimination task. A) Schematic of Luminance Discrimination task. Same as Fig. 1A except for monkeys were trained to select the perceived brighter target. Subthreshold electrical micro-stimulation was applied during the first 500 ms of the 600 ms display period randomly on half of the trials. B) The 14 possible pairs of stimulus conditions. Each luminance level of one target was paired with the middle luminance level of the opposite targets. Which side had the reference and test stimuli was randomized from trial to trial. C) Representative psychometric curves of stimulated (red) and control (black) trials from a single site during an experimental session. Vertical dashed lines show the estimated mu values from the cumulative gaussian fits (see Methods for details). D-E) Population distributions of D) Index of Bias and E) Slope of fit curves in stimulation trials versus non-stimulation trials. The diagonal lines represent unity. The downward-facing triangles denote the mean. The asterisks denote the level of significance (paired t-test, *** for p< 0.001, and ** for p< 0.01). The grey star denotes the example site shown in Fig. S3.

Psychometric curves were compared for stimulation (red) versus non-stimulation (black) trials (grey star highlighted representative site shown in Fig. 5C). Similar to other studies involving perpetual judgements (Moore and Fallah, 2001; Schafer and Moore, 2007), micro-stimulation introduced significant shifts of point of subjective equality (Index of Bias, see Methods), indicating biases of perceptual decisions towards stimulation vectors (paired t-test, N = 60, p = 0.00033, Fig. 5D). In addition, the slopes of fit psychometric curves were significantly steeper when stimulation was applied (paired t-test, N = 60, p = 0.0061, Fig. 5E), suggesting increased discriminability for luminance of target towards stimulation vectors.

Early subthreshold micro-stimulation biased perceptual decisions towards preferred vectors as opposed to stimulation away from the preferred vectors for strategic decisions during comparable stimulation schedule. This suggests a fundamental difference in the role of FEF activation during perceptual versus strategic decision-making.

## Discussion

These experiments demonstrate that FEF plays an active role in choosing strategic saccadic decisions. In a zero-sum mixed-strategy game of Matching pennies, monkeys’ performance approximated the game theoretic equilibrium but with a slight but significant win-stay bias similar to that observed in human and other monkey players (Worthy et al., 2012). Single unit recordings revealed FEF activity were positively correlated with upcoming strategic choices. Specifically, neuronal activity became increasingly predictive of the monkey’s upcoming choices as the time of the saccadic action approached. Artificial perturbation of FEF activity with electrical micro-stimulation influenced the allocation of mixed-strategy choices with the timing and duration of this perturbation being important factors.

Together we take this as strong evidence that FEF plays a sufficient role in saccadic strategic decision process. Below we consider the role of timing in FEF decision formation, a comparison of FEF and downstream SC in these experiments and speculate on how it might bridge the historical biasing signals to the trial-wise action selection.

### Comparisons between SC and FEF during strategic decision-making

One of the major advantages of our experimental design was the strong continuity from the work of Thevarajah et al. (2009). We utilized the same strategic task, replicated experimental procedures and one of the monkeys (monkey B) was used for collecting data in both set of experiments across brain areas.

The FEF sends controls signals to the SC through a dual pathway: an excitatory, direct projection to the intermediate layers of SC (Wurtz et al., 2001), and an indirect pathway that disinhibits inhibitory signals of the substantia nigra through caudate nucleus (Hikosaka et al., 2000). Together a particular FEF vector promotes the corresponding vector in downstream SC through excitatory and disinhibitory means. Indeed, both in FEF and SC preparatory activity gradually ramped up and became increasingly selective throughout warning period of strategic decision-making (Fig. 6A). In addition, subthreshold micro-stimulation significantly biased strategic saccades (Fig. 6B) and strategic selections were biased to preferred direction under most of the perturbation circumstances for both brain areas (Fig. 4). These demonstrated strong functional links between FEF and SC, which may bridge between non-spatial cortical decision variables and trial-wise action selection signal in SC.

**Figure 6.**
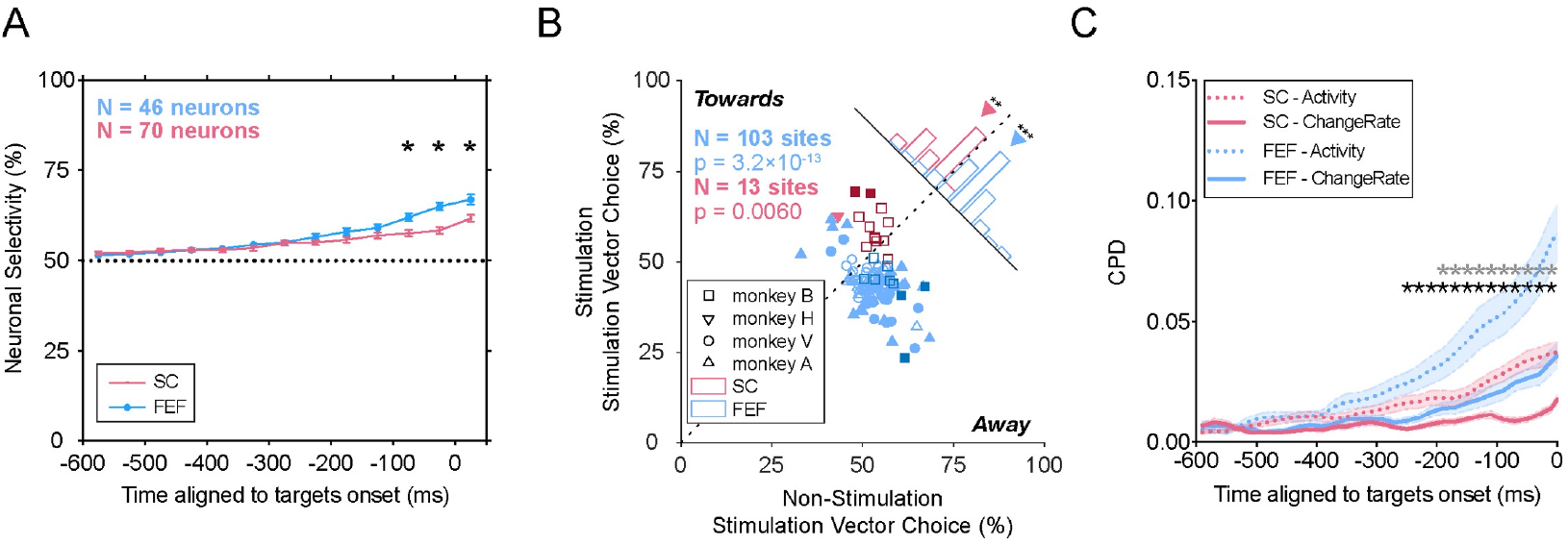
Comparison between FEF and SC activities during Mixed-Strategy task. All SC data obtained from Thevarajah et al. (2009). A) Evolution of neuronal selectivity over time. ROC analysis for FEF (blue) and SC (red) activities. Error bars represent ±SEM. Filled dots indicate significant differences from chance (dashed line) (permutation test, 1000 permutations, p< 0.05). Asterisks denote significant differences between brain areas (two-sample t-test, p< 0.05). B) Differential biasing effect on FEF (blue) and SC (red) when micro-stimulation was applied during the first 500ms of the warning period. Darker squares highlight data from monkey B, on whom both the FEF and SC neurophysiological experiments were conducted, otherwise same as Fig. 3B. C) Evolution of the coefficient of partial determination (CPD) of choice for neuronal activity (dashed) and changing rate of activity (solid). Shaded regions represent ±SEM. Black asterisks denote significant different CPD of changing rate between brain areas, grey for CPD of firing rate (two-sample t-test, p< 0.05).

Yet important differences exist during strategic decision-making between these two brain areas. When stimulating the first 500 ms of the warning period, strategic choices were biased away from and towards the preferred vectors in FEF and SC, respectively (Fig. 6B). Moreover, the changing rate of FEF neuronal discharge was predictive of upcoming strategic choices in a manner independent from the current rate of neuronal activity (Fig.6C, dashed lines) in the FEF (Fig. 6C, solid blue) but not for the SC (Fig. 6C, solid red).

Below we rule out potential explanations for these differences between FEF and SC that could be attributed to confounding factors, and then propose a conceptual framework of our results.

### Ruling out technical confounding factors

Here we outline potential factors that could result in the differential results we observe for FEF and SC during strategic decision-making. Subthreshold micro-stimulation might preferentially activate neurons with ipsilaterally preferred vectors (Bruce et al., 1985), or inhibitory interneurons rather than pyramidal neurons (Lowe and Schall, 2018), or might activate axons leading to postsynaptic responses of negative feedback specifically (Nowak and Bullier, 1998). Alternatively, stimulation might have introduced more disturbance than activation due to the set of stimulation parameters (Histed et al., 2009; McIntyre and Grill, 2000; Murasugi et al., 1993; Nowak and Bullier, 1998; Rattay, 1999). However, none of these attempts to explain why subthreshold micro-stimulation biases choices away from the FEF response fields through technical factors are not applicable to our experiments. The simple fact that supra-threshold stimulation reliably triggered saccades to a contralateral vector and early subthreshold stimulation biased choices towards this same contralateral vector in both the perceptual task and using extended stimulation parameters in the strategic task, rules out these technical explanations.

### Toy model of attractor landscape

Below we introduce a schematic loosely based on attractor dynamics models of decision-making (Wang, 2002; Wong and Wang, 2006). These schematics were inspired by previous work of mechanistic attractor models designed to generate a binary choice (Wang, 2002) and capture the mechanism of time integration (Wong and Wang, 2006) during perceptual decision-making. We admit that this toy schematic is speculative, nevertheless it may provide a useful conceptual framework that captures many of our recording and stimulation results and may provide a useful foundation for testing unresolved questions of FEF mechanisms during strategic decision-making.

The toy schematic depicts the energy level associated with competing decision states as uneven surfaces across time. The ultimate attractors associated with the two alternate choices are represented by the lowest energy basins at the right of each panel (Fig. 7). Here, timing is an orthogonal unidirectional dimension along which energy of decision states steadily descend. The trajectory of the ball running on the landscape is analogous to the decision process in a trial. The trajectories are probabilistic across trials and the illustrations represent conjectures on average.

**Figure 7.**
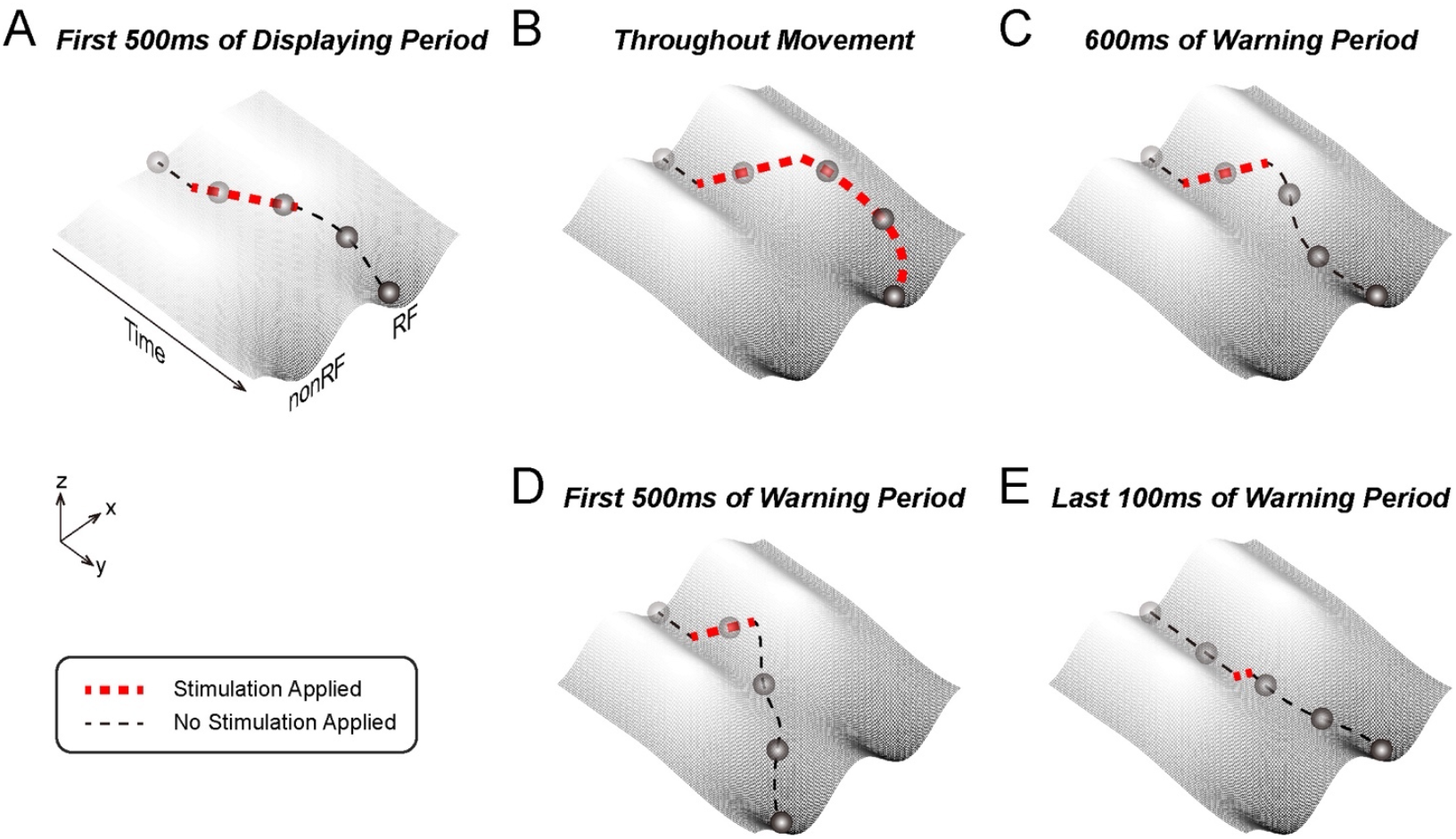
A speculative toy attractor dynamics model to conceptualize our experimental results. A) The perceptual decision process and B-E) strategic decision process during B) stimulation throughout movement, C) 600 ms stimulation, D) 500 ms stimulation, E) stimulation in last 100 ms. Landscapes show energy of decision states. Basins of landscapes denote decision attractors. The ball indicates the decision state. X-axis represents the decision variable, y-axis represents time, and z-axis represents the energy of decision states. The darkening of the ball highlights the progress through time. Black dashed lines are the trajectories of the decision state evolution. Red dashed line denotes the periods during which micro-stimulation was applied in each task.

Our results in the perceptual task (Fig. 7A) are consistent with previous experiments and attractor dynamic models (Wong & Wang, 2006). With equally luminant targets, choices are distributed evenly between the two options because noisy signals effectively nudge the ball stochastically between one attractor or the other. During unequal luminance, the difference in sensory evidence provides a perturbator force increasing the likelihood of choosing the brighter targets. Therefore, perceptual judgements are formed, on average, by the difference in favor of one set of external sensory evidence over another. The addition of electrical micro-stimulation (Fig. 5) introduces an artificial perturbation of the ball towards the direction coded by the site of stimulation, thus increasing the probability of choosing the associated option.

Our stimulation results during the strategic tasks did not follow this traditional schematic for binary choice. With a relatively shallow undulating landscape, perturbation of any duration would nudge the ball towards the stimulation site’s preferred direction on average. Hence, we offer that the landscape during strategic decision-making is more complex. For conceptual purposes, we have shown an early undulating landscape and with a saddle shape late in the warning period that may resolve our observations and capture more strategic and probabilistic decision-making (Fig. 7B-E).

Rather than any external perceptual evidence that can exert influence on attractor dynamics, during strategic decisions it is primarily noise and historical influences (e.g., win-stay bias) that determines the outcome of the competition. In this regard, the symmetric landscape enables a fairly stochastic outcome at the equilibrium probability of 50% with a small trial-wise historical bias (Fig. 1D). That the neuronal activities became more selective over the course of a trial (Fig. 2H) is reflected in diverging trajectories across the landscape as the trial progresses.

The undulating landscape (Fig. 7B-E) of the decision process could help explain the different outcomes when micro-stimulation was applied at various timing schedules in the strategic task. When stimulation was applied during the entire trial (Fig. 7B), a constant extra force is added to the ball, contributing to it finishing towards the preferred attractor. With shorter stimulation, however, the outcome of micro-stimulation became more complicated (Fig. 7CD). The ball is initially perturbed towards the preferred direction, but once stimulation is terminated, the ball rolls down the hill with such momentum that it could continue across the saddle midpoint towards the opposing valley (Fig. 7D). For this particular scenario to occur, the onset and particularly the offset of the stimulation timing would be important. Simulation beyond 500 ms would extend the ball past the saddle and the point of no return (Fig. 7C) whereas only 100 ms of stimulation at the end of the warning period (Fig. 7E) would not provide enough potential energy to jump across the saddle once stimulation was released. In summary, we propose that a more undulating landscape of the strategic decision space could account for the effects of stimulation duration and timing observed in our experiments.

In support of a late saddle on the landscape, not only the representation of current firing rate (location), but also the differential of firing rate (velocity) was predictive of final choices. This is in line with our finding that changing rate of FEF neuronal discharge was predictive of upcoming strategic choices besides the firing rate (Fig. 6C). Since the differential of firing rate (velocity) in the SC was not contributive to strategic choice, we suggest a less undulating landscape such that stimulation applied at any time would result in bias towards the stimulation vector (Fig. 6B).

### Outlook

One of the unexpected findings of this paper was a biasing away effect when stimulation was applied to the FEF early in the warning period. Our work is in line with previous studies suggesting cortical premotor regions may function to similar mechanisms as perturbation also appear to inhibit decision formation or action planning in a manner that interacts with timing (Churchland and Shenoy, 2007; Cunnington et al., 1996). We offered a toy model of an attractor landscape that could account for the effects of stimulation timing and duration on choice direction but this model requires more empirical evidence. First, a more systematic application of stimulation schedules during the Mixed-Strategy task would provide a more detailed view of the FEF decision landscape during strategic decision-making. Second, optogenetic techniques could provide valuable information about the role of specific neuronal sub-types and the direct/indirect pathways from FEF to SC in strategic decision formation. Third, applying micro-stimulation during games with payoff matrices as controlled variable might test the toy model. For instance, in a game whose equilibrium is choosing the preferred vector at a 70% rate, micro-stimulation may bias choices towards preferred direction. But if the reversed biasing effect of stimulation was due to more innate functional properties, stimulation in other game setting might introduce reversed bias anyway.

## Materials and methods

### General methodology

We conducted neurophysiological experiments on three male adult rhesus monkeys weighing approximately 10 kg each. Experiments on monkey B were conducted at Queen’s University in Kingston, Ontario, Canada. Experiments on monkeys V and A were conducted at the Center for Excellence in Brain Science and Intelligence Technology. All respective procedures for experiments were approved by the Queen’s University Animal Care Committee that complied with the guidelines of the Canadian Council on Animal Care and the Animal Care Committee of Shanghai Institutes for Biological Science, Chinese Academy of Science (Shanghai, China), respectively. We strived to replicate identical experimental procedures at both institutions with any significant differences outlined below.

Physiological recordings techniques as well as the surgical procedures have been described previously (Munoz and Istvan, 1998).

The neurophysiology recording chambers were placed over the left frontal eye field with the help of a neuroanatomical atlas (Paxinos et. al., 1999) and structural MRI (Yin et al., 2019) (3T, TR = 2500 ms; TE = 3.12 ms; inversion time = 1100 ms; flip angle = 9°; acquisition voxel size = 0.5 mm × 0.5 mm × 0.5 mm; 144 sagittal slices) scans for each monkey. For monkey B, an upright chamber was placed over a craniotomy at (A23, L20). The chambers for monkey V and monkey A were angled 25° leftward of vertical, targeted towards (A18, L16, D30) and (A24, L15, D30), respectively. The reconstructed axonometric MRI views through the chambers of target neurophysiological sites are shown in Fig. 2A.

All neurophysiological experiments were conducted on Monkey V. Only micro-stimulation experiments were collected on monkey A. For monkey B, neurophysiology recording and blocks of MSstim500 data (see Overview of all experimental blocks) were collected.

### Hardware

Visual stimuli were presented to the monkeys by rear projection (BenQ: TS819ST) onto a tangent screen (monkey B) or by displaying on a monitor screen (monkey V, A – Samsung: DB55E). The screens spanned ±40° horizontal and ±30° vertical of the central visual field. Monkeys’ eye position was recorded by an EYELINK1000 infrared eye tracker (SR Research) at 1000 Hz. Stimuli presentation, data collection, and delivery of reward were real-time controlled by Gramalkn Ryklin Software (monkey B) or by MonkeyLogic (monkey V, A – http://www.monkeylogic.net/). The extracellular single neuron activities were recorded with tungsten electrodes (Frederick Haer; 100K-1M) and the same electrodes were also used for micro-stimulation experiments. Electrodes were lowered by a hydraulic microdrive (Narishige – Monkey B) or a motorized microdrive (NAN Instruments – Monkey V, A).

### Behavioral Tasks

The task design of Mixed-Strategy task, Unpredictable task and Predictable task and stimulation schematics in MSstim500 blocks were identical to that reported in Thevarajah et al. (2009).

#### Mixed-Strategy Task (MS)

To probe the role of FEF in mixed-strategy decision-making, we designed a saccadic version of simplest mixed-strategy game – Matching Pennies (vonNeumann and Morgenstern, 1944).

Subjects competed against a dynamic computer opponent (see Fig. 1A). Subjects were required to maintain central gaze fixation throughout the 800 ms presentation of the fixation point and after its removal during a fixed 600 ms warning period. The fixed warning period and known target locations facilitated advanced selection and preparation of saccades (Dorris and Munoz, 1998). Subjects were free to saccade towards either of two simultaneously presented targets, one of which was presented at the response field of single or multi neurons (RF, preferred), and the other at the mirror-image in the contralateral visual field (non-RF, non-preferred). After fixating the target stimulus for 300 ms, a red box, which indicated the computer opponent’s choice, appeared around one of the targets for 500 ms.

The monkey received a 200 μL liquid reward if both players’ choices matched and nothing otherwise. The computer opponent performed statistical analyses on the subject’s history of previous choices and payoffs and exploited systematic biases in their choice strategy (for specific details, see algorithms 2 from Lee et al. (2004)).

Thus, although monkeys were free to choose, their total payoff was maximized by allocating their choices stochastically between the two targets from trial to trial. On average, monkeys completed 288 ±3 (mean ±SEM) trials per session in this task.

#### Unpredictable Task

The Unpredictable task (Fig. S1A) was identical to the Mixed-Strategy task with two exceptions. First, only a single saccade target was presented on each trial. This target was equally likely to be presented at the preferred or non-preferred direction of the neuron under study. Second, reward was equally likely to be presented or withheld for successful completion of each trial. The square was only shown around the target at the rewarded trials. Therefore, the overall patterns of both choices and rewards were similar for both the Mixed-Strategy and Unpredictable tasks but saccadic choice was under voluntary control in the former case and under sensory instruction in the latter. On average, monkeys completed 183 ±8 (mean ±SEM) trials per session in this task.

#### Predictable Task

The Predictable task (Fig. S1B) was also identical to the Mixed-Strategy task except that a single saccadic target was presented on each trial always at the same location. During one block, the target was always presented in the preferred direction of the neuron, and during a second block, the target was always presented in the non-preferred direction of the neuron. To produce an overall reward rate that was comparable with the other tasks, we opted to give reward on all trials but its magnitude was halved. Therefore, both saccade direction and reward delivery were maximally certain in this condition. Another option, which more closely resembled the reward schedule of the other tasks, would have been to give the regular magnitude reward but on only one-half of the trials. We opted against this latter reward schedule, because it would weaken the rule that matching brings reward (necessary in MS task) when the only possibly matching target presented could not bring reward in all circumstances. On average, monkeys completed 100 ±4 (mean ±SEM) trials for each preferred and non-preferred target in this task.

#### Luminance Discrimination Task (LD)

In the Luminance Discrimination task (Fig. 5A), we kept as many of the parameters the same as possible. Rather than make a voluntary mixed-strategy decision, here monkeys were required to make a perceptual judgement as to which of two visual stimuli were brighter. We replaced the warning period with a 600 ms of visual display period. Two targets were displayed on the screen, while monkeys were required to maintain fixation on the central fixation point. After the fixation point disappeared, monkeys directed a saccade to the perceived brighter target. Monkeys received liquid reward of 200 μL on every correct trial.

We chose 7 luminance levels with equal log step (22.20 cd/m^2^, 26.05 cd/m^2^, 30.57 cd/m^2^, 35.87 cd/m^2^, 42.10 cd/m^2^, 49.40 cd/m^2^, 57.97 cd/m^2^). Luminance of target stimuli were chosen with multiple levels of luminance paired to the median level (Fig. 5B). All pairs of stimuli were presented pseudo-randomly to guarantee 10-15 repeats for each condition. On average, monkeys completed 306 ±6 (mean ±SEM) trials per session in this task.

### Scaling target size with brain magnification factor

To keep discriminability of targets in the Luminance Discrimination task approximately equal across eccentricities of FEF neuron response fields, we used a brain magnification factor (adapted from (Essen et al., 1984); (Strasburger et al., 2011)) to scale the size of the targets with eccentricity:

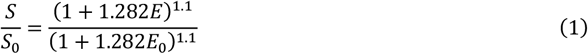

S represents for the radius of targets; E represents the eccentricity of the targets. The formula was designed to maintain discriminability from eccentricities proportional to the setting of S_0_ and E_0_. Hence, arbitrarily, S_0_ was set to be 0.8° and E_0_ was set to be 6°. For simplicity, S was fixed to 0.8° for all targets with E less than 6°.

### Subthreshold micro-stimulation

To determine whether direct activation of FEF could bias monkeys’ strategic and perceptual choices, we applied micro-stimulation during the Mixed-Strategy and Luminance Discrimination tasks, respectively. The methodology closely followed that of Thevarajah et al. (2009). The threshold of an FEF site was established using micro-stimulation (100 ms duration, 300 Hz, and 0.25 ms biphasic pulses) in a pure fixation paradigm. Stimulation threshold was defined as the minimum current intensity required to trigger saccades towards the stimulation vector of the site on approximately 50% trials. From this site-specific threshold, we established subthreshold micro-stimulation parameters by decreasing stimulation frequency from 300 Hz to 100 Hz and reducing current intensity, if necessary, such that saccades to the point that saccades were never elicited during the warning or discrimination periods. The timing and duration during which subthreshold micro-stimulation was applied for different schedules is outlined in Figure 3A. In all experiments, micro-stimulation was applied randomly on half of the trials in a session. Current amplitude averaged 28±1.2 μA (mean ±SEM) in Mixed-Strategy task and 23 ±1.6 μA (mean ±SEM) in Luminance Discrimination task.

### Overview of all experimental blocks

On a typical experimental session, blocks were conducted in the following order: Mixed-Strategy, Unpredictable and finally Predictable tasks resulting in more data from the former than latter blocks. Stimulation experiments were generally conducted on separate days from recording experiments and usually at sites with multi-unit activity because the activity of a single neuron could not be isolated. We strived to run as many of the stimulation conditions on one site as possible to permit within site comparisons across timing schedules and with the Luminance Discrimination task (see Fig. 4D and Fig. S4). The Venn diagram in Fig. S2 illustrated all the experimental blocks and their intersections.

### Data analysis

Trials were aborted online under any of the following conditions. 1) Monkeys’ eye position exceeded the 3-degree-radius check window around the fixation point. 2) After target presentation, monkeys’ eye position left the fixation window either faster than 70 ms (i.e., anticipatory saccade) or slower than 350 ms (i.e., slow response). 3) Monkeys failed to hold fixation on targets for 300 ms on the newly acquired target. The first 20 trials from each block were discarded from analysis offline to allow subjects time to adjust to the new task conditions.

Offline data analysis was performed using MATLAB, version R2017b (MathWorks) and customized computer programs.

#### Neuronal Inclusion

The Delayed Saccade task was performed before neuronal recording and stimulation blocks aiming to characterize neuronal properties and neurons’ response fields (Zhang et al., 2021).

Neurons satisfying all the criteria below were included for further analyses. 1) Contralateral response field (in the right visual field) characterized with Delayed Saccade task. 2) Either must be classified as a visual neuron, or a delay neuron, or a motor neuron in Delay Saccade task, as defined by criteria in (McPeek and Keller, 2002). 3) During Mixed-Strategy task, the activity in the 50 ms after targets presentation must be significantly greater than that in the first 100 ms after fixation point offset (paired t-test, α = 0.05).

#### Neuronal Selectivity

We used signal detection theory (Swets et al., 1978) to determine how well an ideal observer of FEF activity could predict which choice the monkey would make. The separation between the distributions of activity for the preferred and non-preferred trials of the neuron was estimated from the area under receiver-operating characteristic (ROC) curves (Britten et al., 1996; Newsome et al., 1989), which is indicated as neuronal selectivity.

#### Single Trial Activity Approximation

We approximated single trial neuronal activity using a causal kernel, where the approximation of the firing rate at time t that depends only on spikes fired before t was calculated using a window function that vanishes when its argument is negative. The window function w was defined as follows (Dayan and Abbott, 2001):

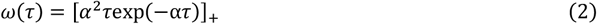

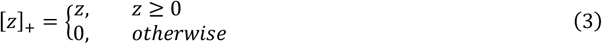

where τ represents time relative to the timing window, 1/α determines the temporal resolution of the resulting firing rate estimate, and the notation [z]_+_ for any quantity z stands for the half-wave rectification operation. We approximate firing rate with temporal resolution 1/α = 30 ms.

#### Analysis of contribution of changing rate of neuronal discharge to strategic choices

To examine the effect size of changing rate of neuronal discharge to the strategic choices, in regard to the accountabilities of neuronal firing rate per se, we computed the coefficient of the partial determination (CPD). CPD quantifies the proportion of the variance accounted for by a set of independent variables. For the independent variables X_2_, CPD is defined as follows (Abe and Lee, 2011):

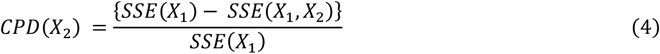

where SSE(X) stands for the sum of squared errors in a regression model that include X. Thus, the CPD indicates the extra fraction of variance that can be accounted for by including the variables X_2_ in the model that already includes X_1_.

We computed the extra fraction of variance accounted by both *changing rate of neural discharge* and *neuronal firing rate*, respectively.

### Stimulation effect

To measure whether micro-stimulation biased monkeys’ strategic choices, we compared choice proportions to RF target for stimulation trials and non-stimulation trials for all experimental sites. We used chi-square statistic as a single number to evaluate the stimulation effect on strategic choices (Fig. 4D). Negative numbers of chi-square statistic were taken if the biasing effect was away from the preferred vector.

To measure whether micro-stimulation biased monkeys’ choices during the Luminance Discrimination task, we fit monkeys’ psychometric functions of the proportion of choices directed towards the neuron’s RF target versus the log of luminance of that RF target with a cumulative gaussian curve. We extracted the Index of Bias and the slope to describe this function:

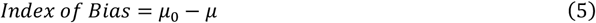

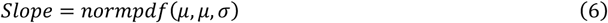

μ_0_ is a constant, representing the theoretical points of subjective equality, which equals the log of the median luminance. μ and σ are parameters for expectation and standard deviation of the fit gaussian curve. Separate psychometric functions were constructed for stimulation and non-stimulation conditions. Indexes in two conditions were compared for population sites with paired t-test (α = 0.05).

#### Saccade Reaction Time (SRT)

We defined the saccade onset time to be the earliest time point when the velocity of eye movement surpassed 30 degrees per second. Monkeys’ saccade reaction time (SRT), was defined as the interval between the go signal and the onset of the saccade. For the Mixed-Strategy, Unpredictable and Predictable tasks, the go signal was the onset of targets and, and for the Luminance Discrimination task, the go signal was the offset of the fixation point.

To estimate how much the stimulation variable can explain the variance in saccade reaction time (SRT), in regard to the choice influences on SRT, we fit a general linear model of SRT as function of stimulation and choice and the interaction term in sessions with micro-stimulation:

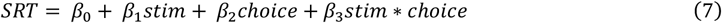

where *stim* and *choice* denote dummy variables (*stim* = 1 when stimulation applied, and 0 otherwise; *choice* = 1 when RF target was chosen, and 0 otherwise). We took *β*_*1*_ +*β*_*3*_ as the additional SRT of stimulation when RF target was chosen, and *β*_*1*_ as the additional SRT of stimulation when non-RF target was chosen (Fig. S5).

## Acknowledgements

This work was supported by grants from the CAS Hundreds of Talents Program. We thank the members of the Dorris lab and members of Gu lab for their help in all phases of the study and Y. Wang for constructive feedback regarding the manuscript.

## Author Contributions

S.X. and M.C.D. conceived the project and designed the experiments. S.X. performed the experiments on monkey V and monkey A and analyzed all experimental data. A.A. collected the data from monkey B. Y.G. devoted conceptual contribution to the manuscript. M.Y. supported with algorithm adaptation for the computer opponent. X.W. dedicated on experimental setting up. J.T. helped with animal care. D.T. collected data in brain area SC. S.X. drafted manuscript. S.X. and M.C.D. edited and revised manuscript.

## Supplementary figures

**Figure S1.**
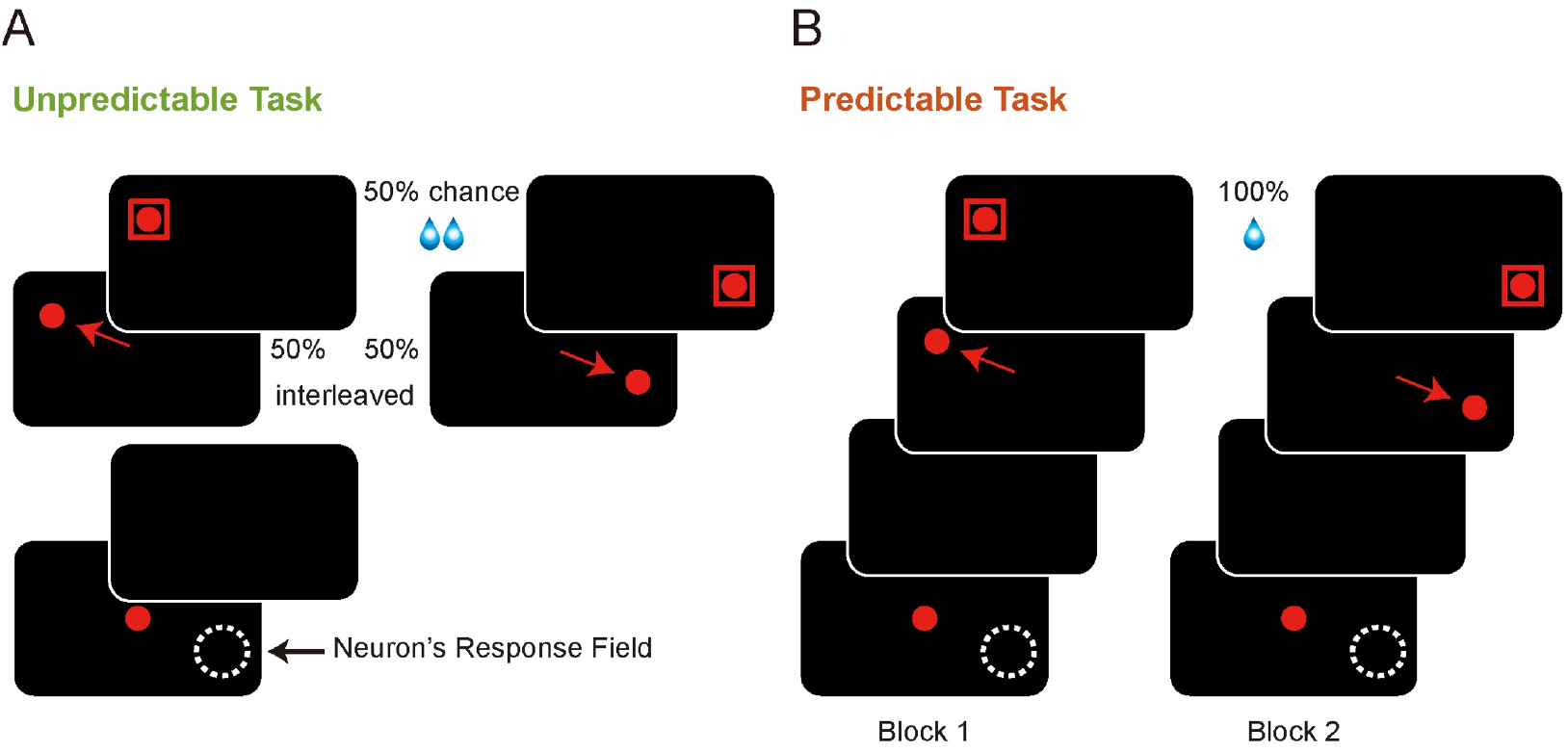
Schematics of control tasks. The white dashed circle represents the response field of a neuron for a given site of FEF. For both of the tasks, the monkeys were required to make a saccade to the single target that was presented on each trial to receive a liquid reward. A) In the Unpredictable task, target presentation was randomized equally between the RF and mirror-image location. Reward was dispensed at a 50% rate for successful completion of trial. The red square mimicked the choice of the computer opponent and was only presented during rewarded trials. B) The Predictable task was conducted in two blocks. In one block, the target was always presented in the RF, and in the other block, always at the mirror-image position. Reward was dispensed on every completed trial but at half the volume of other tasks.

**Figure S2.**
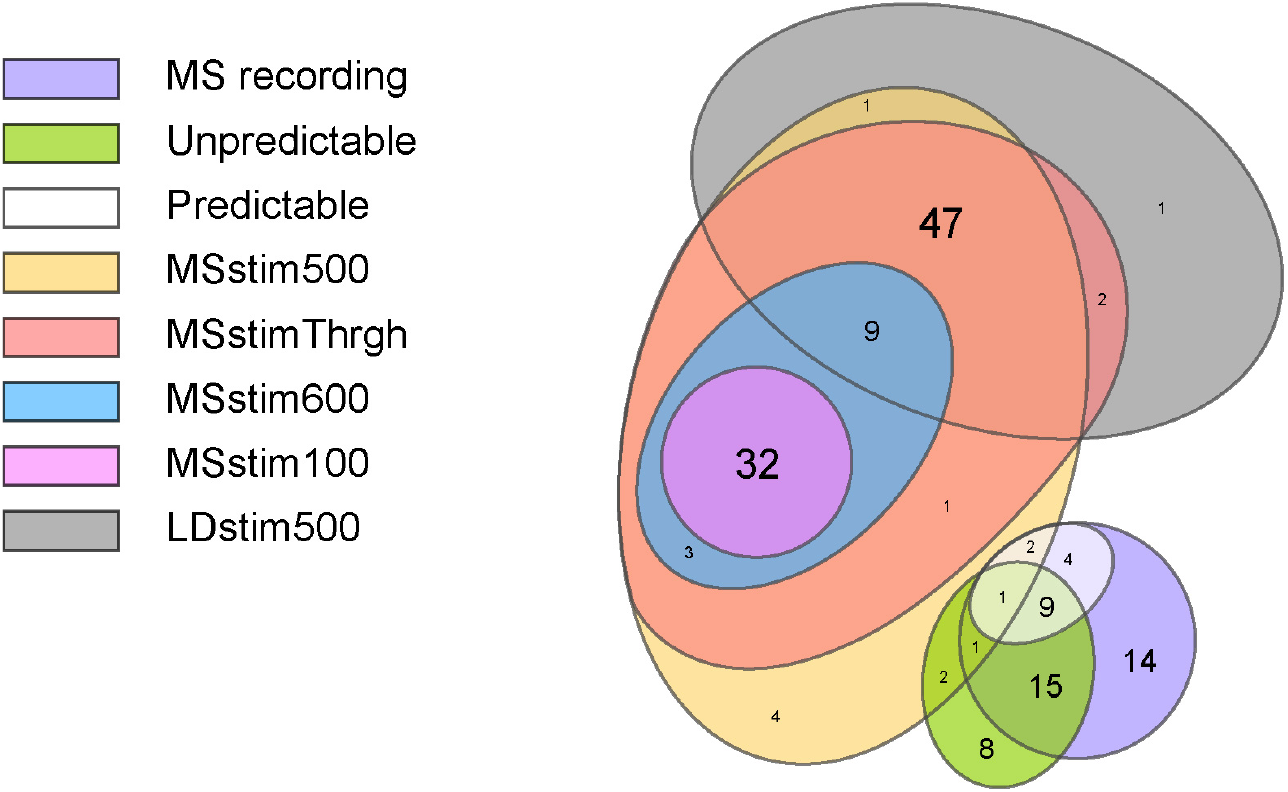
Overview of all experimental blocks. In this Venn diagram, shared areas denote experiments conducted on the same monkeys, neural sites and experimental sessions. Ovals represent sets of experimental blocks. Colors denote different experimental conditions. Numbers of intersecting blocks shown for each irregular-shape intersection.

**Figure S3.**
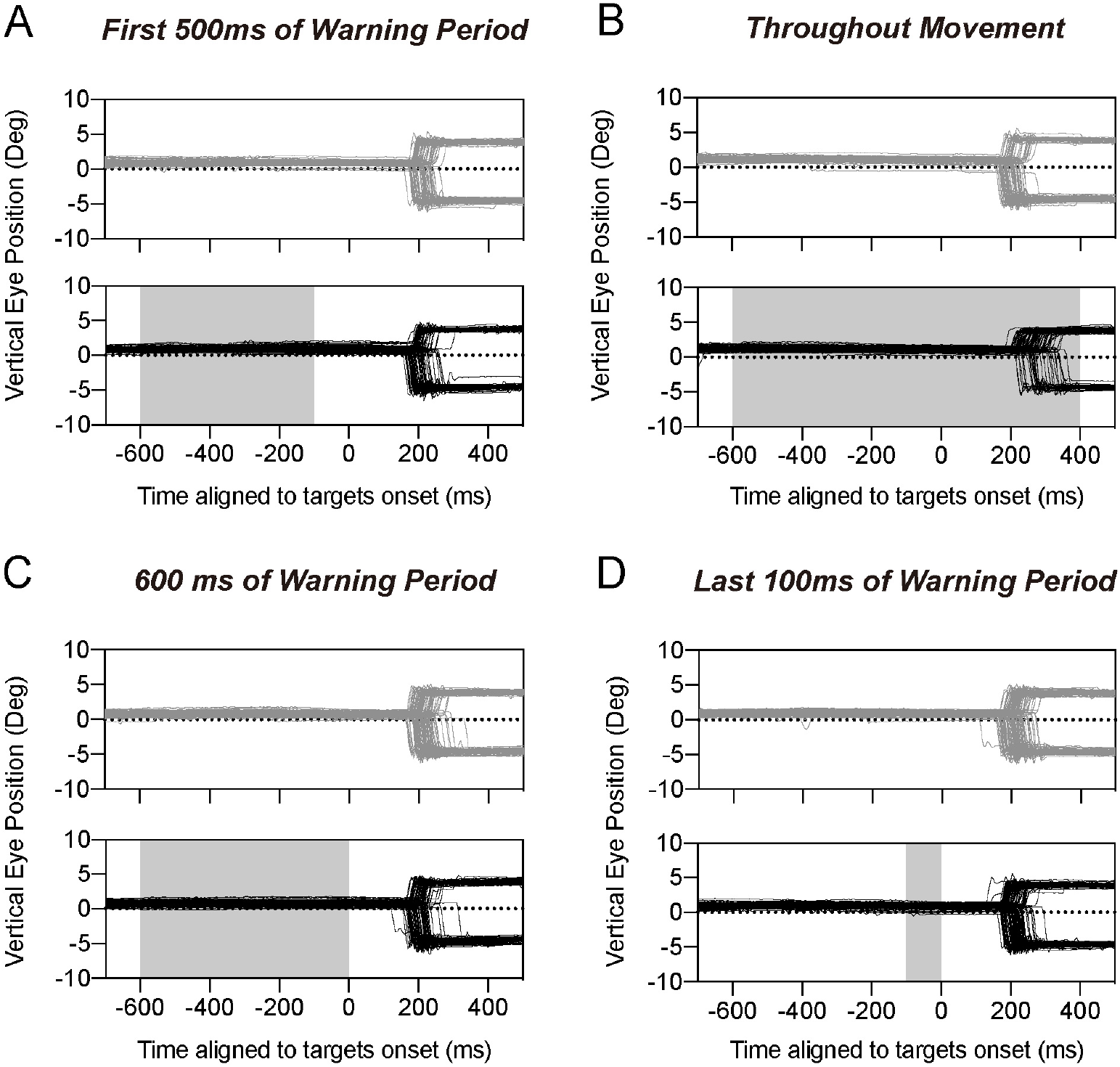
Effect of subthreshold electrical micro-stimulation on eye position. Eye traces of an example site. Vertical eye position in time. Upward deflections denote movements towards the RF target and downward deflections denote movements towards non-RF target when stimulation was applied during the A) first 500 ms of Warning Period, B) throughout movement, C) 600 ms of Warning Period, and D) last 100 ms of Warning Period. Grey traces (top panels) denote non-stimulation conditions and black traces (bottom panels) denote stimulation conditions. The shaded grey areas indicate the period during which micro-stimulation was applied.

**Figure S4.**
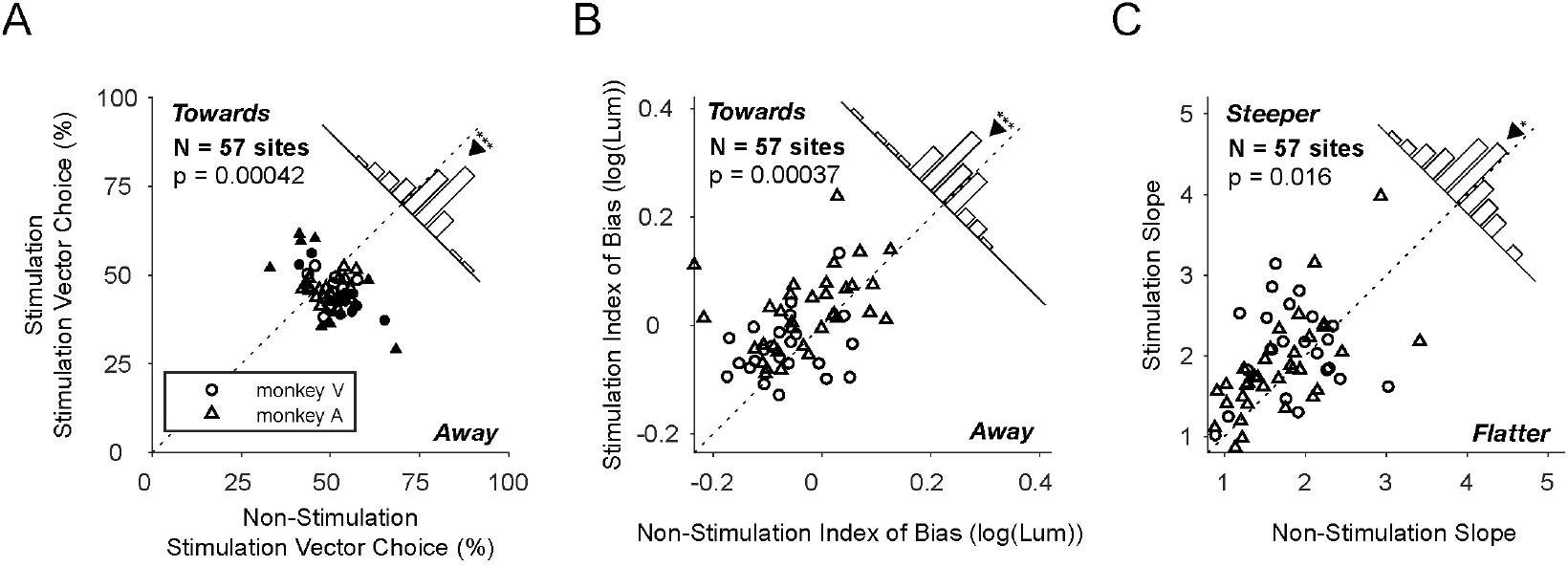
Effects of stimulation applied for 500 ms of warning/displaying period at sites where both Mixed-Strategy and Luminance Discrimination task were conducted (N = 57 sites). A) Format same as Fig. 3B. B and C) Format same as Fig. 5D and Fig. 5E.

**Figure S5.**
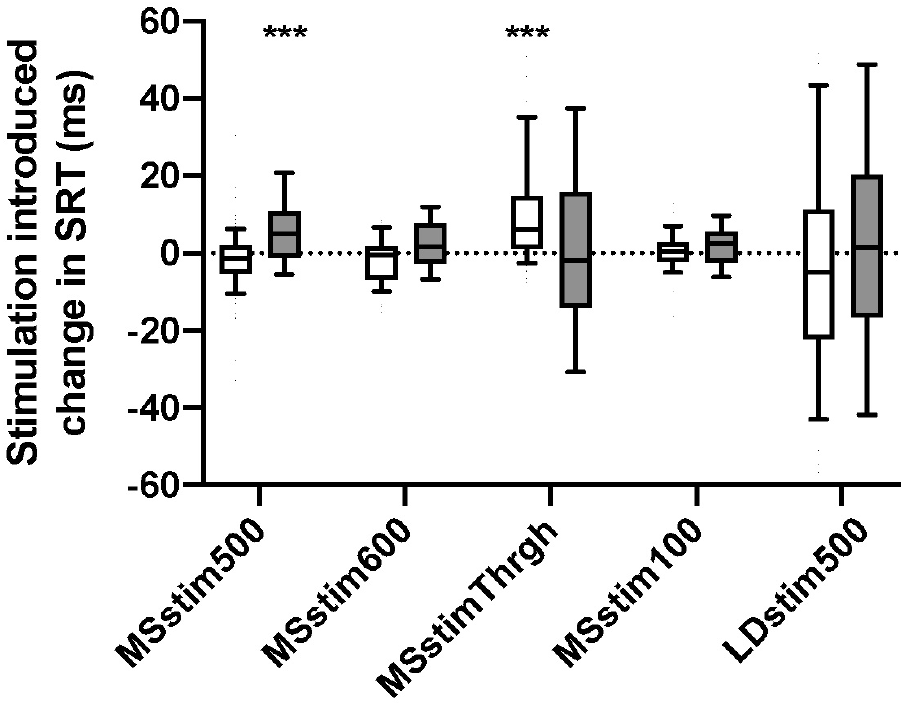
Effects of stimulation on saccade reaction time (SRT) across tasks. Changes in SRT introduced by stimulation for non-RF choices (white) and RF choices (grey) are shown in box plots. Horizontal dashed lines represent medians. Boxes indicate percentiles ranging from 25%-75%. Extending whiskers show percentiles ranging from 10%-90%. The horizontal dashed line denotes no change. Asterisks show level of significance (Two-sided sign test on medians against zero, *** represents p< 0.001).

